# Interstitial and cerebrospinal fluid exchanging process revealed by phase alternate labeling with null recovery MRI

**DOI:** 10.1101/2021.07.26.453795

**Authors:** Anna M. Li, Jiadi Xu

## Abstract

**Purpose:** To develop Phase Alternate LAbeling with Null recovery (PALAN) MRI methods for the quantification of interstitial to cerebrospinal fluid flow (ICF) and cerebrospinal to interstitial fluid flow (CIF) in the brain.

**Method:** In both T_1_-PALAN and apparent diffusion coefficient (ADC)-PALAN MRI methods, the cerebrospinal fluid (CSF) signal was nulled, while the residual interstitial fluid (ISF) was labeled by alternating the phase of pulses. ICF was extracted from the difference between the recovery curves of CSF with and without labeling. Similarly, CIF was measured by the T_2_-PALAN MRI method by labeling CSF, which took advance of the significant T_2_ difference between CSF and parenchyma.

**Results:** Both T_1_-PALAN and ADC-PALAN observed a rapid occurrence of ICF at 67±56 ms and 13±2 ms interstitial fluid transit times, respectively. ICF signal peaked at 1.5 s for both methods. ICF was 1153±270 ml/100ml/min with T_1_-PALAN in the third and lateral ventricles, which was higher than 891±60 ml/100ml/min obtained by ADC-PALAN. The results of the T_2_-PALAN suggested the ISF exchanging from ependymal layer to the parenchyma was extremely slow.

**Conclusion:** The PALAN methods are suitable tools to study ISF and CSF flow kinetics in the brain.

## Introduction

Cerebrospinal fluid (CSF) plays an essential role in maintaining the homeostasis of the central nervous system, providing buoyancy to the brain (1), serving as an important route for the removal of a variety of waste products produced by cellular metabolism (2). Its circulation has been the subject of speculation and experiment for more than one hundred years. However, its formation and circulation are still under debate. (3–9) Conventionally, it is believed that the anterior choroidal arterial blood secretes CSF via choroid plexuses (CP) inside the brain ventricles (80–90%), and CSF flows unidirectionally along subarachnoid spaces to be absorbed into venous sinuses. (3,10–12) Some other pieces of evidence supported that CSF continuously exchanges with the interstitial fluid (ISF) in its surrounding brain parenchyma, which depends on hydrostatic and osmotic forces, i.e. the transependymal flow. (13–17) Notably, the recent rediscovery of the brain lymphatic system, dubbed as the glymphatic system, suggested a portion of the subarachnoid CSF recirculates through the brain parenchyma and exchanges with the ISF and then flow back to CSF (18–21). The glymphatic system has drawn intensive attention since its discovery because it appeared to clear off soluble amyloid-beta (Aβ) from the brain parenchyma (22–25). Consequently, it is crucial to develop a clinically useful prognostic tool for ISF-CSF exchange measurement that can be used for the diagnosis and evaluation of brain diseases. Many non-invasion MRI methods, diffusion-based method (26,27), and the time-of-flight-based MRI method (28) were implemented to examine the bulk flow inside the ventricles and glymphatic vessels. However, examining CSF and ISF exchanging process still relied on MRI contrast agents such as the intra-cranial (19,29–32) or intrathecal injection (33–35) of gadolinium-based contrast agents and intravenous D-glucose infusion. (36,37) Contrast agent-based MRI methods are far from ideal for routine and repeated measurements on patients.

The T_1_, T_2_, and apparent diffusion coefficient (ADC) are significantly different between CSF (T_1_=3.0 s, T_2_= 300 ms at 11.7T; ADC=3 μm^2^/ms) and ISF (T_1_=1.8 s, T_2_= 40 ms at 11.7T; ADC=0.7 μm^2^/ms). (36,38) In principle, T_1_, T_2_, or ADC can selectively label either CSF or ISF to monitor the ISF-CSF exchange process. However, in practice, it is a challenge to label CSF without attenuate ISF and vice versa. In the current study, we proposed a novel MRI strategy, dubbed as Phase Alternate LAbeling with Null recovery (PALAN), that null one of the components (ISF or CSF) and label the other component (CSF or ISF) by flipping the phase of pulses. The ISF-CSF exchanging process shaped the recovery curve of CSF or ISF. A strategy closely resembles the flow-sensitive alternating inversion recovery arterial spin labeling (ASL) method. (39) ICF measured via T_1_ and ADC difference and CIF measured via T_2_ difference are named T_1_-PALAN, ADC-PALAN, and T_2_-PALAN, respectively. This series of quantitative PALAN methods provide novel tools to further understand CSF and ISF exchanging processes in the brain.

## Methods

### MRI experiments

The study was carried out under the approval of Johns Hopkins University animal care and use. For this study, ten female mice (C57BL/6J) aged 11-12 months were purchased from Jackson laboratory. All MRI experiments were performed on a horizontal bore 11.7 T Bruker Biospec system (Bruker, Ettlingen, Germany). A 72 mm quadrature volume resonator and a 2×2 mouse phased array coil was used as a transmitter and receiver. All animals were anesthetized using 2% isoflurane in medical air, followed by 1% to 1.5% isoflurane for maintenance during the MRI scan. The respiratory rate was monitored via a pressure sensor (SAII, Stony Brook, NY, USA) and maintained at 70-90 breaths per minute. The B_0_ field over the image slice was adjusted using field mapping and second-order shimming.

### MRI Pulse Sequences

Fig. 1a,b, and c show the schematic diagrams of the T_1_, ADC, and T_2_-based PALAN sequences. For the T_1_-PALAN method, a hyperbolic secant (HS) inversion pulse (15 ms) was applied. At the CSF null inversion time (*TI*_null, CSF_=2 s), the CSF signal was at the zero baseline while part of the ISF signal had recovered. One Z-filter composited by two 90-degree pulses (Gaussian pulse with 1.4 ms width) was applied at the *TI*_null, CSF_. The phase of the second 90-degree pulse in the Z-filter was alternated by 180-degree, flipping the parenchyma longitudinal magnetization up (control) and down (label). CSF recovery during eleven post-labeling delays (PLDs) (0, 0.1, 0.2, 0.4, 0.6, 1, 1.5, 2, 3, 4, 5 s) were recorded. The recovery from the labeling pulse was different from the original recovery curve due to the presence of ICF. Four pairs of control/label parenchyma signals were collected to measure the labeling efficiency, and eight pairs of CSF signals were recorded for the CIF measurement. A turbo spin-echo (TSE) MRI with long echo time (LE-TSE) was implemented to read the CSF signal and to suppress the parenchyma signal with TE = 245 ms, pre-scan delay 5 s, TSE factor = 96, slice thickness = 1 mm, a matrix size of 96 × 96, and a resolution of 0.17× 0.17 mm^2^.

**Figure 1.**
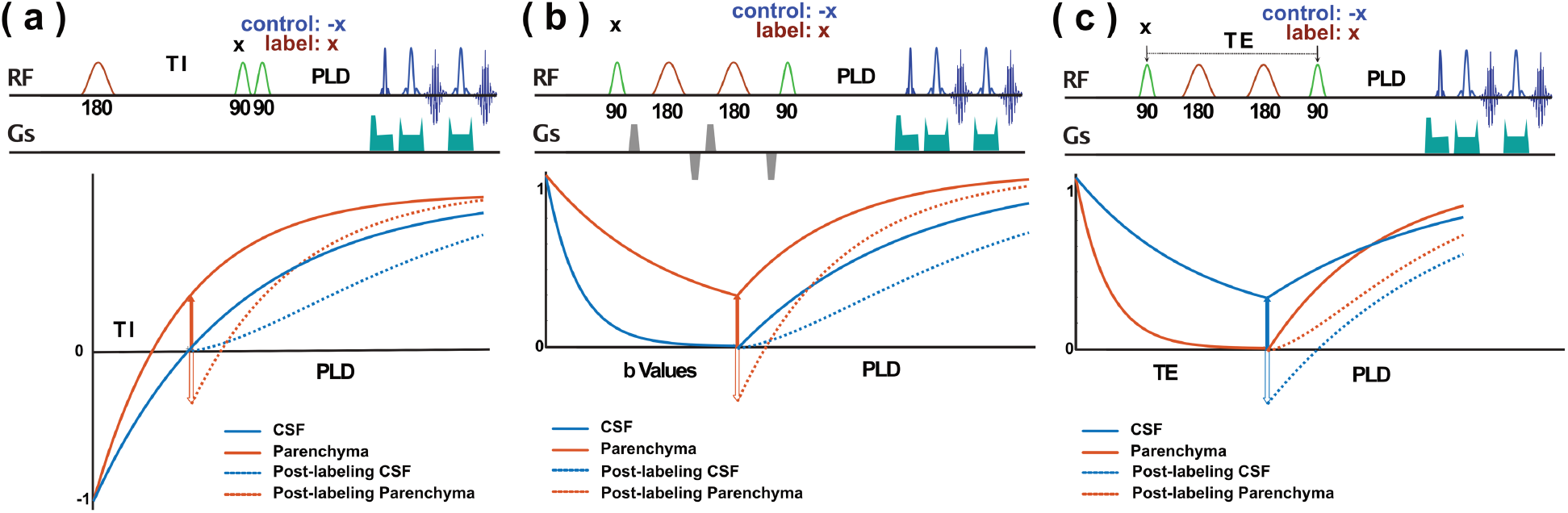
(a) Illustration of the T_1_-based phase alternate labeling with null recovery (T_1_-PALAN) sequence. The signal changes of CSF and parenchyma without labeling are solid lines, after labeling are dashed lines. After the inverting pulse, at the CSF null time, a pair of 90-degree pulses, i.e., Z-filter, is apply. The phase of the second 90-degree pulse in the Z-filter is alternated and can flip the parenchyma longitudinal magnetization up (control) and down (label). The difference between the dashed and the solid CSF recovery curves is due to the ISF-CSF exchange. A TSE MRI with long echo time (LE-TSE) is implemented to readout the CSF signal and to suppress the parenchyma signal. (b) Illustration of the apparent diffusion coefficient-based phase alternate labeling with null recovery (ADC-PALAN) sequence. The signal changes of CSF and parenchyma without labeling are solid lines, after labeling are dashed lines. An improved motion-sensitized driven-equilibrium (iMSDE) module is applied to suppress the CSF signal with high b values, while strong parenchyma signal is preserved. The phase of the second 90-degree pulse in the iMSDE is alternated to flip the parenchyma longitudinal magnetization up (control) and down (label). Similar to T_1_-PALAN, the ISF-CSF exchanging process introduces the difference in CSF recovery curves and LE-TSE is used for the CSF imaging. (c) Illustration of the T_2_-based phase alternate labeling with null recovery (T_2_-PALAN) method. The signal changes of CSF and parenchyma without labeling are solid lines, after labeling are dashed lines. Like the ADC-PALAN, a Carr-Purcell-Meiboom-Gill (CPMG) module is applied to null the ISF signal. The recovery curve of ISF is modulated by the flow from CSF to ISF.

ADC-PALAN method, much like the T_1_-PALAN method, can also extract ICF. Instead of applying one T_1_ preparation module, an improved motion-sensitized driven-equilibrium (iMSDE) module (40) was applied to suppress the CSF signal with high b values (2100 s/mm^2^), but left some residual parenchyma signal. In our study, the total iMSDE module was 30 ms, the gradient length was 3 ms. Mao pulses (41) with 6 ms width was used for the 180-degree pules and Gaussian pulses (1.4 ms width) were used for the two 90-degree pulses in the iMSDE module. The phase of the second 90-degree pulse in the iMSDE module was alternated by 180-degree to obtain the control and label images.

In the T_1_-PALAN and ADC-PALAN multi-PLD studies, a single axis slice was collected at −0.7 mm from the anterior commissure (AC) to cover the left ventricles (LV) and third ventricle (3V). Common belief states that CP is found in the caudal lateral ventricles. (42,43) To determine CSF contributions from CP to ventricles in T_1_-PALAN and ADC-PALAN methods, we collected two high-resolution single PLD (1.5 s) slices (−0.7 and 0.9 mm from AC). ICF maps that covered the rostral and caudal LV (n=5) were obtained. High-resolution T_2_ weighted images with a long TE turbo spin-echo (LE-TSE) sequence determined the CP locations. The scan time for each LE-TSE experiment was 13 minutes with TR/TE= 6 s / 106 ms, RARE factor=32, slice thickness=0.5 mm, a matrix size of 256 × 256 within a FOV of 16 × 16 mm^2^.

The measurement of CSF backflow from ventricles to parenchyma, i.e., CIF, was achieved with the T_2_-PALAN method. Carr-Purcell-Meiboom-Gill (CPMG) module was used in the T_2_-PALAN to null the parenchyma ISF signal. The CIF process modulated the recovery curve of parenchyma. The label and control images were collected by alternating the phase of the second 90-degree pulse at the end of the CPMG module by 180-degree. TSE with short TE (TE = 5 ms) was used to acquire parenchyma MRI images with a pre-scan delay of 5 s, TSE factor = 16, slice thickness = 1.5 mm, FOV=1.6×1.6 mm^2^. A matrix size of 32× 32 was used for the CIF kinetic curve measurement with four pairs of control/label images, and a matrix size of 64× 64 was used for the high-resolution CIF images with 32 pairs of control/label images. The total experimental time for the high-resolution CIF images was 31 minutes. 9 PLDs (0, 0.5, 1, 1.5, 2, 2.5, 3, 4, and 6 s) were acquired for the CSF labeling efficiency measurement and the CIF buildup curves with the T_2_-PALAN method. In the T_2_-PALAN multi-PLD studies, a single axis slice was collected at −0.7 mm from AC for the CSF optimization, and the slice at −4.1 mm from AC was used for the parenchyma signal optimization. Two slices (−0.7 and −4.1 mm from AC) were collected for the high-resolution single PLD (1.5s) CIF maps.

### Data Analysis

All MRI images were processed using custom-written MATLAB scripts (MathWorks, R2020a). The ICF (Δ*S*_*CSF*_) or CIF signals (Δ*S*_*Parenchyma*_) were calculated by subtracting the label and control images following

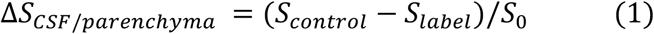

The S_0_ images were collected by setting PLD=10 s for parenchyma and 15s for CSF in all PALAN methods.

The recovery of the parenchyma signal in the T_1_-PALAN and ADC-PALAN after the T_1_ or iMSDE preparation module can be described as

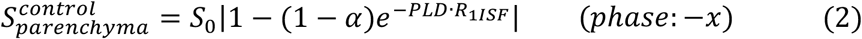

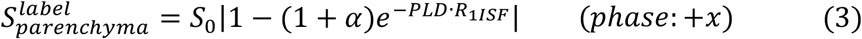

where *α* was the parenchyma signal after the Z-filter, i.e., the labeling efficiency. *R*_1*ISF*_ was the T_1_ relaxation rate of the parenchyma. In practice, nonzero noise background was present in the MRI image due to the magnitude Rician noise. Therefore, the reliable way of extracting the labeling efficiency was fitting the difference image 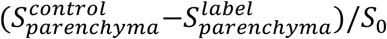 following

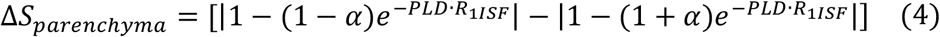

Similarly, the difference between the two CSF recovery curves after the CPMG module in the T_2_-PALAN method was given by

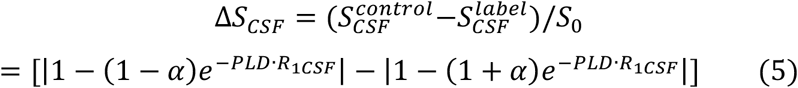

*R*_1*CSF*_ was the CSF T_1_ relaxation rate.

A standard single-compartment kinetic model was applied to quantify the ICF from the observed CSF signal reduction Δ*S*_*CSF*_, assuming instantaneous exchange of labeled spins from parenchyma to ventricles. The observed CSF signal reduction was described by the difference between the sum over the series of delivered magnetization units to CSF from ISF (the arterial input function, AIF) and the clearance of the magnetization by the relaxation of the CSF (the impulse residue function, IRF) (44–46).

The IRF function was given by

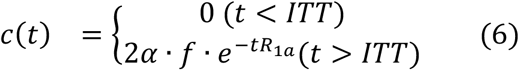

Where *α* was the ISF labeling efficiency of the T_1_ or iMSDE preparation module, f was the ICF flow in units of volume of ISF delivered per volume of CSF per unit time (ml/100ml/min). ITT was the ISF transit time, i.e., the time for ISF to reach CSF after labeling. The *R*_1*a*_ was a labeling decay rate due to the limited bolus duration.

The observed CSF reduction Δ*S* was the convolution of the AIF and the IRF functions, i.e.,

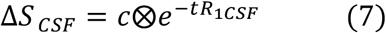

Then,

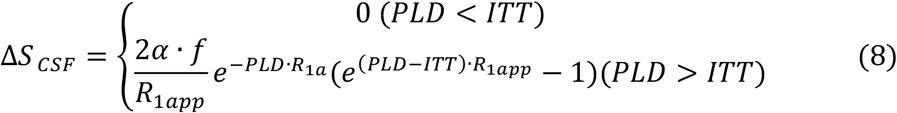

where *R*_1*app*_ = *R*_1*a*_ − *R*_1*CSF*_. When measuring CIF rate by Δ*S*_*Parenchyma*_, the same equation can be used except the *R*_1*app*_ = *R*_1*CSF*_ − *R*_1*ISF*_ and

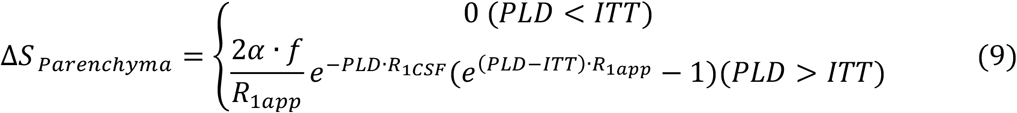

f was the CIF flow in units of volume of CSF delivered per gram ISF per unit time (ml/100mg/min).

A paired-sample t-test was performed for the comparison between CIF values of the rostral and caudal LV using the MATLAB built-in function “*ttest*”. t-test is considered statistically significant for p<0.05 and highly significant when p<0.001.

## Results

### ISF to CSF flow measured by T_1_-PALAN

The typical T_1_-PALAN CSF and parenchyma recovery curves, together with the ICF Δ*S* maps, are shown in Figs. 2(a-d). The post-labeling parenchyma signal curve shows a typical inversion recovery curve. Theoretically, the MRI signals with PLD less than 0.4 s were negative after the labeling pulse as seen from the standard inversion recovery curves, but the usage of magnitude-reconstructed images resulted in positive values. The parenchyma signal difference curves using Eq. 4 yielded the labeling efficiency of α=0.21±0.02. The CSF recovery curves show a clear difference between control and label pulses, indicating high ICF values. The magnitude Rician noise background led to a positive CSF signal at PLD=0 s. The CSF difference curves, i.e., ICF kinetic curves, show a clear buildup and decay pattern that peaks at PLD=1.5 s. Eq.8 fitted perfectly with the curve, which gave ICF = 1153±270 ml/100ml/min, and ITT = 67±56 ms. The ICF ΔS values for the rostral and caudal LV are 0.015±0.013 and 0.034±0.01 (p=0.022, n=5), respectively (Fig. 2e).

**Figure 2.**
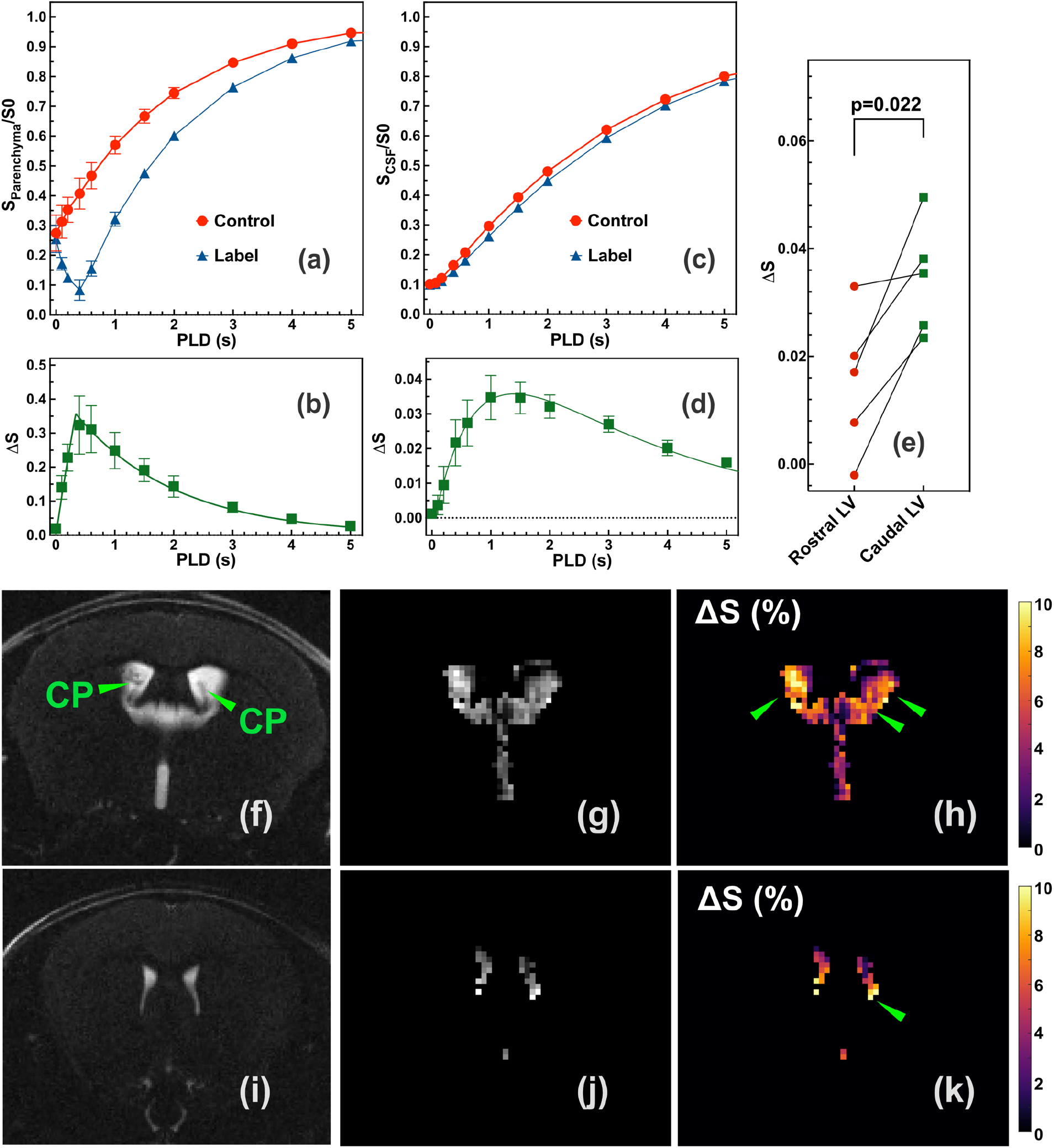
(a) Averaged parenchyma recovery curves for the whole slice post control/label pulse (n=3) as a function of the post labeling time (PLD) for the T_1_ based null recovery with phase alternate labeling (T_1_-PALAN) sequence. (b) The difference between control and label of the parenchyma recovery curves. Solid line is theoretical fitting curves with Eq.4 (R^2^=0.97). (c) Averaged CSF recovery curves for lateral ventricle (LV) and the third ventricle (3V) with the post control/label pulse (n=3) as a function of the PLD for the T_1_-PALAN sequence. (d) The difference of the averaged CSF recovery curves, i.e. the ISF-CSF flow (ICF) kinetic curve, as an function of PLD. Solid line is theoretical fitting curves with Eq.8 (R^2^=0.99). (e) The ICF Δ*S* values for the rostral and caudal LV. (f, i) The high-resolution T_2_ weighted images by LE-TSE for the two slices collected. Choroid plexuses (CP) are indicated with green arrows. (g, j) The typical control images for T_1_-PALAN method. (h, k) The corresponding ICF maps acquired with T_1_-PALAN. The regions with hyperintensity ICF values are indicated with green arrows.

Figs. 2g, h, j, and k are typical T_1_-PALAN control images and ICF maps of the brain ventricles. The high-resolution T_2_ maps shows the CP in the LV (Figs. 2f and i), and the CP is clearly visible in the caudal LV regions. The ICF values were calculated from Eq. 8 with *R*_1*a*_ = 1.4 *s*^−1^, ITT=67 ms, and a labeling efficiency of 0.21. Across the ventricles, the ICF rates were not uniform, and the majority of ISF flowed from the bottoms of the LV as indicated in Figs. 2h and k with green arrows.

### ISF to CSF flow measured ADC-PALAN

We performed the CSF and parenchyma optimization as a function of the PLD for ADC-PALAN shown in Fig. 3. As suggested by the b-dependent ADC measurement on the parenchyma and CSF (supplemental Fig.S1), the CSF signal was less than 1% with b=2100 s/mm^2^ than the signal with b=0. The difference between label and control parenchyma signals yielded a labeling efficiency of 0.033±0.012. Due to the signal reduction introduced by ADC and T_2_, the observed labeling efficiency for parenchyma was significantly lower than those of the T_1_-PALAN method, which led to a much-reduced ICF signal 4.2±0.5 × 10^−3^ (PLD=1.5 s) compared to 35±4.5 × 10^−3^ (PLD=1.5 s) in the T_1_-PALAN method. Eq.8 fitted perfectly with the ICF kinetic curve and gave ICF = 891±60 ml/100ml/min, and ITT = 13±2 ms. The ICF Δ*S* values for the rostral and caudal LV were 3.9±1.9× 10^−3^ and 4.4±1.4× 10^−3^ (p=0.66, n=5), respectively. The much lower signal obtained by the ADC-PALAN method stopped us from obtaining the ICF maps.

**Figure 3.**
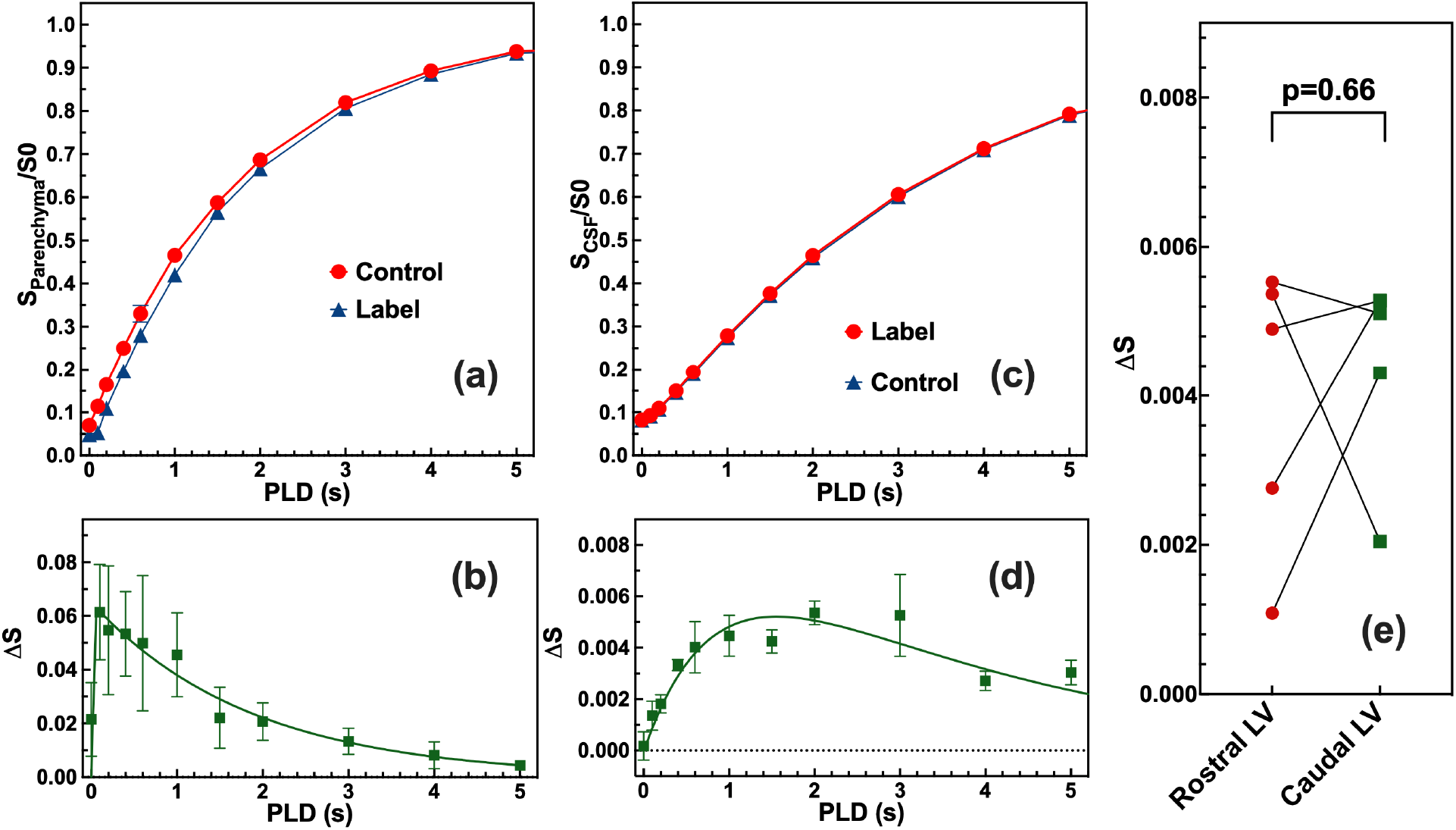
(a) Averaged parenchyma recovery curves for the whole slice after the control/label pulse (n=3) as a function of the post labeling time (PLD) for the apparent diffusion coefficient based null recovery with phase alternate labeling (ADC-PALAN) sequence. (b) The difference of the parenchyma recovery curves. Solid line is the theoretical fitting curve in Eq.4 (R^2^=0.97). (c) Averaged CSF recovery curves with the control/label pulse (n=3) as a function of the PLD for the ADC-PALAN sequence. (d) The difference of the averaged CSF recovery curves, i.e. the ISF-CSF flow (ICF) kinetic curve, as a function of PLD. Solid line is theoretical fitting curves with Eq.8 (R^2^=0.95). (e) The ICF Δ*S* values for the rostral and caudal lateral ventricle (LV).

### CSF to ISF flow measured by T_2_-PALAN MRI

The backflow from CSF to parenchyma i.e., CIF was measured with the T_2_-PALAN method, and the results are presented in Fig. 4. Differed from the ADC and T_1_ PALAN method, the CSF signal was maintained and labeled in the T_2_-PALAN sequence. The CSF recovery difference between the post-labeling and the original curves with Eq. 5 gives the labeling efficiency of 0.28±0.03. The parenchyma recovery curves after the label/control pulse (Fig. 4c) shows a barely noticeable difference, which indicates a negligible flow rate from CSF to the parenchyma. The CIF Δ*S* signal is under 0.22%. (Fig. 4d) To obtain a reliable estimate of the ICF values, we addressed the labeled CSF strong interference and the overall low signal in the map (Fig. 4g) by performing the high-resolution CIF maps with 32 averages at a slice with much fewer ventricles, i.e., −4.1 mm from AC (Figs. 4h-j). The overall CIF ΔS signals for the ROIs shown in Fig. 4h were found to be 0.076±0.027% at PLD=2 s (n=3) since simulation found the maximum ICF signal at 2s. The maximum CIF value was estimated to be 15±6 ml/100mg/min with Eq.9, assuming ITT=0 ms, *R*_1*CSF*_ = 0.33 *s*^−1^ and *R*_1*ISF*_ = 0.56 *s*^−1^.

**Figure 4.**
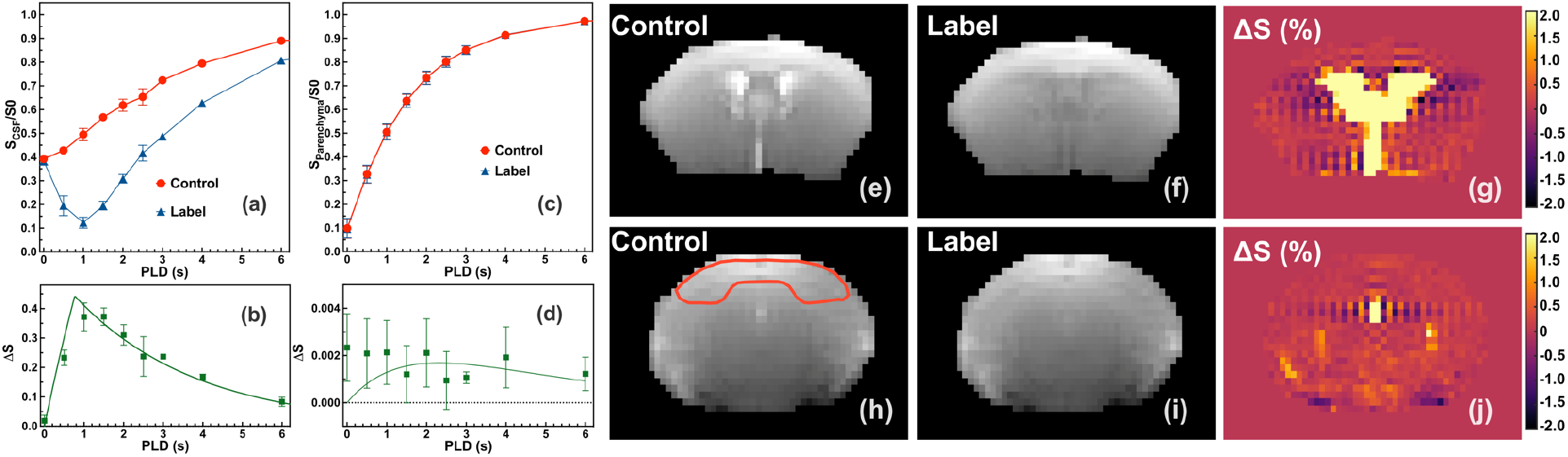
(a) Averaged CSF recovery curves for the CSF in ventricles after the control/label pulse (n=3) as a function of the post labeling time (PLD) for the T_2_ based null recovery with phase alternate labeling (T_2_-PALAN) sequence. (b) The difference of the CSF recovery curves. Solid line is the theoretical fitting curve with Eq.5 (R^2^=0.93). A label efficiency 0.28 was obtained from the fitting. (c) Averaged parenchyma recovery curves with the control/label pulse (n=3) as a function of the PLD for the T_2_-PALAN sequence. (d) The difference of the averaged parenchyma recovery curves (n=3), i.e. the CSF-ISF flow (CIF) kinetic curve, as an function of PLD. Solid line is the simulated curve using Eq.9 by assuming ICF=20 ml/100mg/min, ITT=0 ms, *R*_1*CSF*_ = 0.33 *s*^−1^ and *R*_1*ISF*_ = 0.56 *s*^−1^. The typical control (e, h) and label images (f, i) by the T_2_-PALAN method for two slices. The CIF Δ*S* maps obtained by T_2_-PALAN method following Eq. 9 for PLD=2s (g,j). The typical ROIs used for extracting CIF Δ*S* **v**alues are indicated in (h).

## Discussion

This study proposed a series of non-invasive PALAN MRI methods to measure the ISF and CSF interchange, i.e., ICF and CIF flows, by selectively labeling ISF with T_1_ and ADC contrasts and CSF with T_2_ contrast. The T_1_-PALAN pulse sequence provided much higher signals than the ADC-PALAN. Both T_1_-PALAN and the ADC-PALAN methods suggested that ISF flow into CSF rapidly. On the contrary, CSF to the parenchymal regions far away from ventricles was a pronouncedly slow process and barely observable with the spin labeling method.

T_1_-PALAN has its advantage in terms of obtaining the kinetic curves. In our study, *T*_1a_=1/*R*_1a_ =0.7 s was found, which was far less than the parenchyma T_1_ relaxation time of 1.8 s. Therefore, the bolus duration generated by the T_1_ module was in the order of 0.7 s. In the ADC-PALAN, the blood signal was fully suppressed by the high b value (2100 s/mm^2^) and cannot be labeled. In contrast, the blood signal can fully recover during the T_1_ preparation module due to the blood in-flow effect and the labeling efficiency was close to one for blood. Therefore, the T_1_-PALAN ICF values contained strong signal from the blood through CP, i.e., blood-cerebrospinal fluid barrier arterial spin labeling, (47) and the ICF value (1153±270 ml/100ml/min) was higher than the value measured by the ADC-PALAN (891±60 ml/100ml/min). The contamination from CP in T_1_-PALAN was further validated by the significantly different ICF Δ*S* values of the rostral and caudal LV with the T_1_-PALAN (Fig. 2e). The similar ICF values measured by ADC-PALAN in both rostral and caudal LV confirms that the observed ICF signal mainly came from ISF not blood by ADC-ALAN. (Fig. 3e)

The recent pseudo-continues ASL method has measured the blood-cerebrospinal fluid barrier (BCSFB) flow in mice to be 13-20 ml/100 ml/min, i.e. a total flow of 0.52-0.8 μL/min by assuming LV volume 4 μL. (47) Gd MRI agent is unable to measure the BCSFB flow because of its inability to pass the BCSFB. However, D-glucose penetrates BCSFB with the help of abundant glucose transporter on the BCSFB. Hence dynamic glucose enhancement can be used to assess the BCSFB flow. (36,37) Interestingly, the BCSFB ASL signal in mouse LV was mainly found at the top of both LV and no signal observed in 3V. The ICF hyperintensity regions measured by the T_1_-PALAN method are visible in 3V and at the bottom of the LV. (Figs. 2h and k)

The novel non-invasive PALAN methods proposed here provided an effective tool to reveal the CSF production and circulation in the brain. Current study quantified the flow rate of ICF, i.e., 891-1153 ml/100ml/min in LV and 3V. A recent invasive study on mouse brain states the CSF outflow from both LV and 3V is in the order of magnitude of 0.1 μL/min (48), which is minute compare to the ICF flow obtained by our current study (32-44 μL/min in 4 μL LV). Our result indicated that the CSF secreted from parenchyma is reabsorbed by the periventricular ependymal layers as indicated in Fig. 5b. No fresh CSF is generated in this process. The reason behind rapid turnover of the CSF with the ependymal layer is under speculation. Water continuously moves in and out of the ependymal layer, a process resembles the intracellular and extracellular water movement, is one possible reason for the rapid turnover number. The difference between CSF secreted from CP (0.52-0.8 μL/min) (47) and the total outflow (0.1 μL/min) (48) suggests that a large portion of the CSF in ventricles is absorbed by the surrounding parenchyma. But due to the large mass of the parenchyma, kinetically speaking, the absorption is a prolonged process (less than 15 ml/100mg/min) as confirmed by the T_2_-PALAN measurement. In Fig.4g, the strong CSF signal (~ 37%) obscures CIF signal of the parenchyma adjacent to the ependyma layer (<1%), preventing us from making a solid conclusion that the CIF rate was high in those parenchyma regions. Another force that facilitates the CSF and ependymal layer exchanging process is the pulsatile nature of the CSF flux, which appears to be craniocaudally oriented during cardiac systole and in the reverse direction during diastole (49–51) This model not only explains the extremely high turnover rate of the CSF measured by PALAN method, but also perfectly explains the radioactive-labeled substances study in CSF, in which the infused tracer crosses the periventricular ependymal layer and can be found in the extracellular space. (17,52,53) However, more work is needed to confirm the above hypothesis. One last note, in theory, T_1_-PALAN can also measure the backflow from CSF to parenchyma by setting a null time to parenchyma signal, but the brain white matter and gray matter have different T_1_ null time, making it impossible to null the entire parenchyma at the same time.

**Figure 5.**
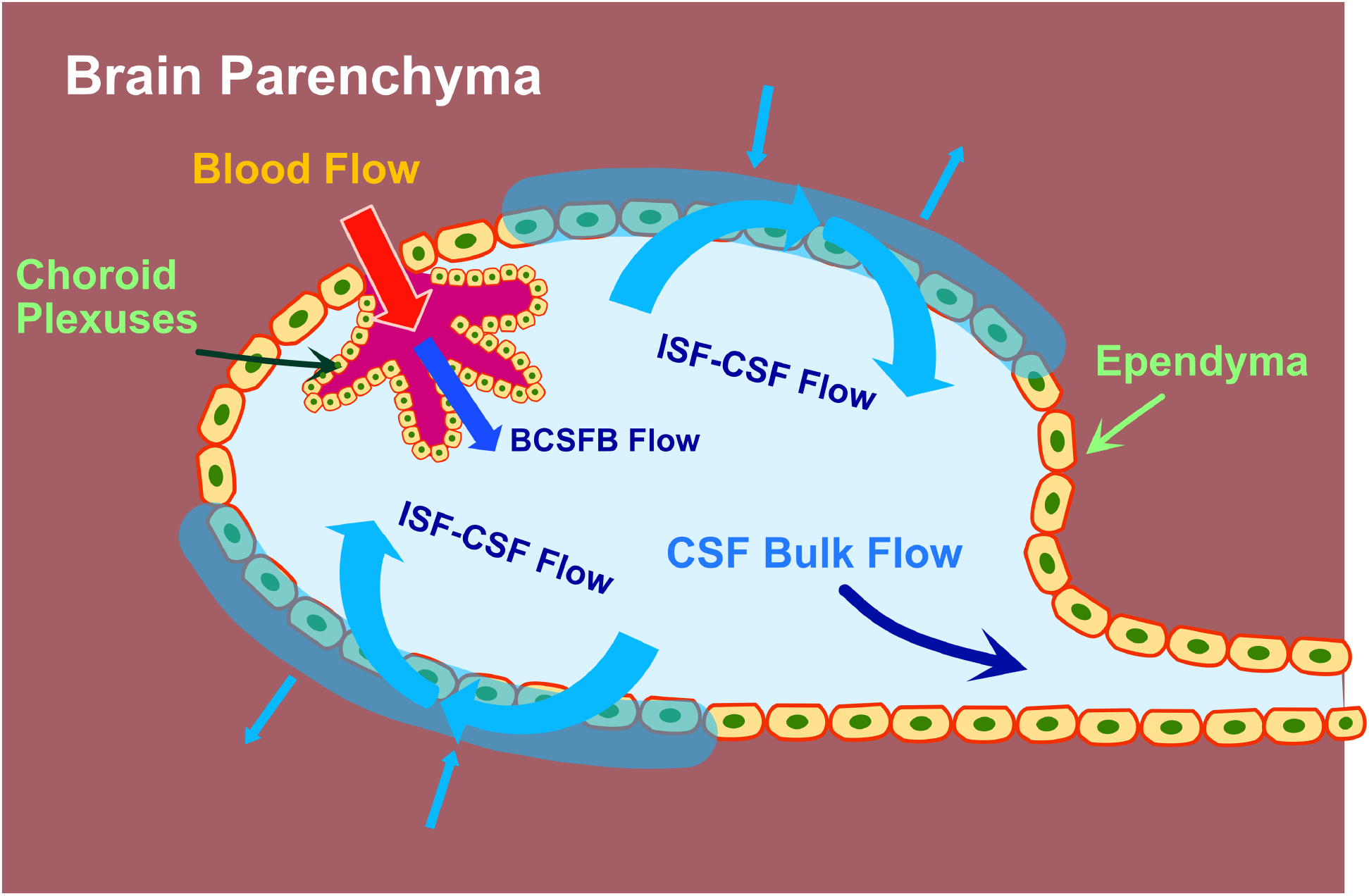
The schematic diagram of the CSF flow in ventricles revealed by the PALAN methods. CSF is primarily secreted from the choroid plexuses (CP) provided by the anterior choroidal arterial blood. There is a rapid exchange between the interstitial fluid (ISF) in the ependymal layer and the CSF in the ventricles. However, the exchanging process from the ependymal layer to the regions far away from ependyma is an extremely slow process. Eventually, CSF flows out of ventricles and exits into the subarachnoid space (SAS) at the cisterna magna.

In summary, PALAN MRI methods proposed here provide technically achievable tools to examine the association between ICF and functional decline in many neurodegeneration diseases. They complement the invasive MRI method with agents and give a complete view of the ISF and CSF exchanging process.

## Conclusion

We presented T_1_, T_2_, and ADC PALAN schemes as new translational MRI methods to quantify in vivo ISF-CSF and CSF-ISF exchange processes in the mouse brain. PALAN offered ICF images with high sensitivity and quality. We quantified the ICF flow rate from the parenchyma to the ventricles with the T_1_-PALAN and the ADC-PALAN methods, which was consistent with the values measured by the MISL method. The T_1_-PALAN suggested the ISF exchange from ependymal layer to the parenchyma is one lagging process and is challenging to be measured by the spin labeling method.

